# Evaluation of Genomic and Proteomic Expression of Surfactant Protein D in the Mouse Retina

**DOI:** 10.1101/2020.11.20.391078

**Authors:** Frederico Vieira, Johannes W. Kung, Faizah Bhatti

**Author notes:** **Author Contributions:** F.V. (first author) - Conception and design, acquisition, analysis and interpretation of data, drafting and revision of the article, final approval J.W.K. - Design, acquisition, analysis, and interpretation of data, drafting the article, final approval F.B.- Conception and design, analysis and interpretation of data, drafting and revision of the article, final approval. **Corresponding author:** Faizah Bhatti, M.D., M.S. The Children’s Hospital at The University of Oklahoma Health Sciences Center, 1200 Children’s Ave. 7^th^ Floor North Pavilion, Oklahoma City, OK 73104.

## Abstract

Surfactant Protein D (SP-D), an essential protein related to innate immunity, is expressed in multiple tissue types throughout the body. A closely-related protein, Surfactant Protein A (SP-A), is present in the mouse retina and is associated with neovascularization (NV) in the Oxygen-Induced Retinopathy (OIR) mouse model, mimicking retinopathy of prematurity (ROP). We hypothesized that SP-D would be present in the retina and is also associated with OIR and ROP, which is one of the leading causes of pediatric blindness due to increasing survival rates of extremely preterm newborns. In our study, we did not detect SP-D in the mouse retina through proteomic and genomic investigation at baseline and in pathways known to up-regulate SP-D in other mammal tissues. It is therefore unlikely that SP-D participates in neovascularization in the mouse retina.

## Introduction

Retinopathy of prematurity (ROP) is a disease that is seen in premature newborns (1) and is related to abnormal development of the retinal vasculature and neovascularization *ex utero*. Worldwide, ROP is a leading cause of acquired childhood visual impairment and blindness. The etiology of this complex disease is still not completely understood. While exposure to high levels of oxygen and nutritional status are important in the pathology of ROP (2), inflammation has also been shown to be a key risk factor (3).

Surfactant Protein-D (SP-D), as the name implies, was first described in the surfactant substance of the lung (4). SP-D, along with Surfactant Protein A (SP-A), are recognized as collectin proteins. The collectin family of proteins shares homology in their basic monomeric structure, which is comprised of collagen-like regions attached to non-collagenous domains (5). The collagen domain and neck domain are essential in maintaining the intermediary tetramer protein structure (6). The non-collagen N-terminal domain is a cysteine-rich region critical for oligomerization occurring through disulfide bridging necessary for the mature structure of the protein (7). The last non-collagen domain is the carbohydrate recognition domain (CRD), or lectin domain. This domain is what defines the function of all collectin proteins, since they can bind to carbohydrate and lipid epitopes of different sizes and shapes from multiple microorganisms when polymerized in their tridimensional structure (8, 9). The CRD binds to multiple eukaryotic receptors necessary for innate immunity, including Toll-like receptors (TLR) 2, TLR-4, and CD14 (10, 11).

Preterm infants have developmental immaturity of their adaptive immune response (12), as well as pathology related to a deficiency of surfactant and surfactant proteins. Recent investigations have found that these proteins are not exclusive to the lungs and are present in multiple organ systems throughout the body (13). We have previously reported the expression of SP-A in the mouse retina and showed that it can be expressed in cultured human Müller glial cells by ligand-receptor activation of toll-like receptors (TLR) TLR-2 and 4. We also demonstrated the up-regulation of SP-A in the murine model of oxygen-induced retinopathy (OIR). Furthermore, animals with gene targeting of SP-A had a significant reduction in neovascularization (14).

SP-A and SP-D and are both present on chromosome 10 in humans and are co-expressed in lung tissue, sharing significant homology in structure and function. Therefore, we hypothesized that SP-D is present in the mouse retina in a pattern similar to that of SP-A. Furthermore, we hypothesized that SP-D is up-regulated by exposure to oxygen stress, as well as activation of TLR-2 and TLR-4. To study this, we used a variety of protein and RNA expression assays to identify and quantify SP-D. These experiments were performed in mouse retinal tissue at baseline and after receptor activation using the toll-like receptor (TLR) ligands 2 and 4. Expression was also measured in the mouse OIR model. After we found no protein or RNA expression, we utilized mass spectrometry to confirm our results. Finally, OIR was issued in SP-D null mice (SP-D ^−/−^ or knockout [KO]) to demonstrate that the absence of SP-D does not affect the phenotype of mouse retina exposed to hyperoxia and inflammatory stress.

## Materials and Methods

### ANIMALS

All animal procedures adhered to the American Association for Laboratory Animal Science (AALAS) guidelines and the Association for Research in Vision and Ophthalmology (ARVO) Statement for the Use of Animals in Ophthalmic and Vision Research, and were approved by the University of Oklahoma Health Sciences Center Institutional Animal Care and Use Committee. All mice were exposed to a standard 12-hour light/dark cycle and fed standard mouse chow. C57BL/6J wild-type (WT) mice were obtained from Jackson Laboratory (Bar Harbor, ME, USA). Surfactant Protein D null mice (SP-D ^−/−^) were a kind gift from the Korfhagen Lab in the Division of Pulmonary Biology from Children’s Hospital Research Foundation (Cincinnati, OH, USA). The generation and background of these mice have been reported previously in detail (15). PCR was used to determine the genotype of all pups prior to analysis (16). After confirmation of genotype, wild-type mice were euthanized, and their retinas were collected at postnatal day 0 (P0), P2, P5, P7, and P14, and in adulthood (ages varied from P48 to 6 months of age). Mice at P0 and P2 were deeply anesthetized via cryoanesthesia and decapitated. Mice at P5 and older were euthanized by CO_2_ asphyxiation. Tissue was then harvested for each experiment as described below.

### GENOTYPING

PCR confirmed the genotypes of animals used in experiments. DNA samples were obtained by a tail fragment or ear punch digestion with 50 mL 50 mM NaOH digest buffer for 1 h at 95 °C. The pH was adjusted to 7.0 by titration with 7 μL buffer containing 100 μM Tris (HCl) and 10 μM EDTA. DNA was added to Quick-Load *Taq 2x* Master Mix (New England Biolabs, Ipswich, MA, USA) and respective primers.

For SP-D detection in WT animals, the forward primer was generated in the first intron and the complimentary reverse primer was developed for the second exon, which includes the codon for translation start with the sequence. The primers for the detection of the knockout gene (KO) were developed based on the mouse created by Korfhagen *et al.* by the integration of a vector neo nucleotide sequence. The neo sequence deletes the initiating methionine and translation initiation sequence ATG from the second exon (15, 16). For the rd1 mutation on the Pde6b gene, primers were used according to the methods of Blazek et al. (17). All primers were purchased from Eurofins Genomics (Louisville, KY, USA). Sequences are described in Table 1.

**Table 1.**
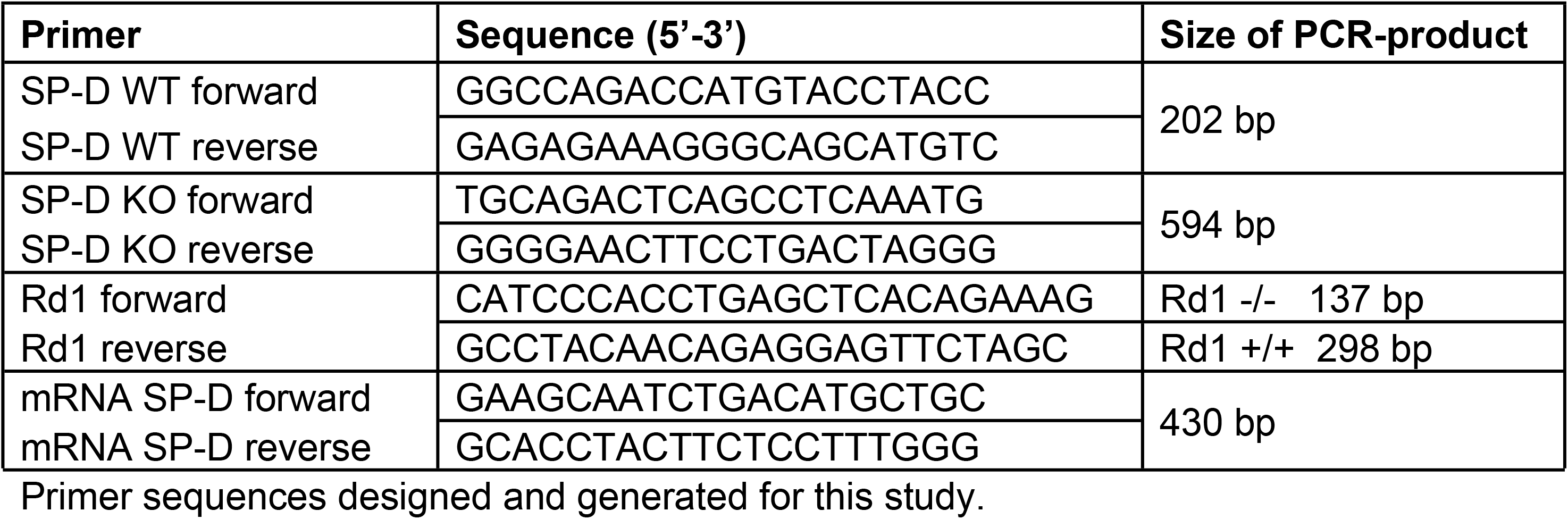
List of primer sequences.

### LOCALIZATION AND EXPRESSION OF SP-D IN MOUSE RETINA

Our first objective was to determine whether SP-D mRNA and protein were expressed by retinal tissue during various developmental time points in the WT mouse. Measurements in SP-D ^−/−^ mice were used as a control. Localization and expression were evaluated by PCR for mRNA, by immunohistochemistry (IHC) for protein localization, and by western blotting and ELISA for protein measurements.

#### Detection of SP-D-encoding mRNA

RNA was extracted from mouse whole lung and whole retina using PureZOL RNA reagent from BIO-RAD Laboratories (Hercules, CA, USA) with the method described by Chomczynski *et al.* (18). RNA samples were tested for purity by Nanodrop 2000 from Thermo Scientific (Lenexa, KS, USA). cDNA was synthesized from mRNA using the qScript cDNA synthesis kit from Quanta Biosciences (Beverly, MA, USA) and used for PCR at 61 °C with OneTaq Quick-Load 2x Master Mix from New England Biolabs (Ipswich, MA, USA) and primers specifically binding to cDNA derived from mature mRNA, but not to genomic DNA. The forward primer targets the end of exon 1, skips intron 1, and ends in exon 2, which includes the ATG sequence for the start codon for protein translation (Figure 1). The reverse complementary primer starts at the end of exon 2, skips intron 2, and ends in exon 3, generating a 430-bp fragment (Table 1).

**Figure 1.**
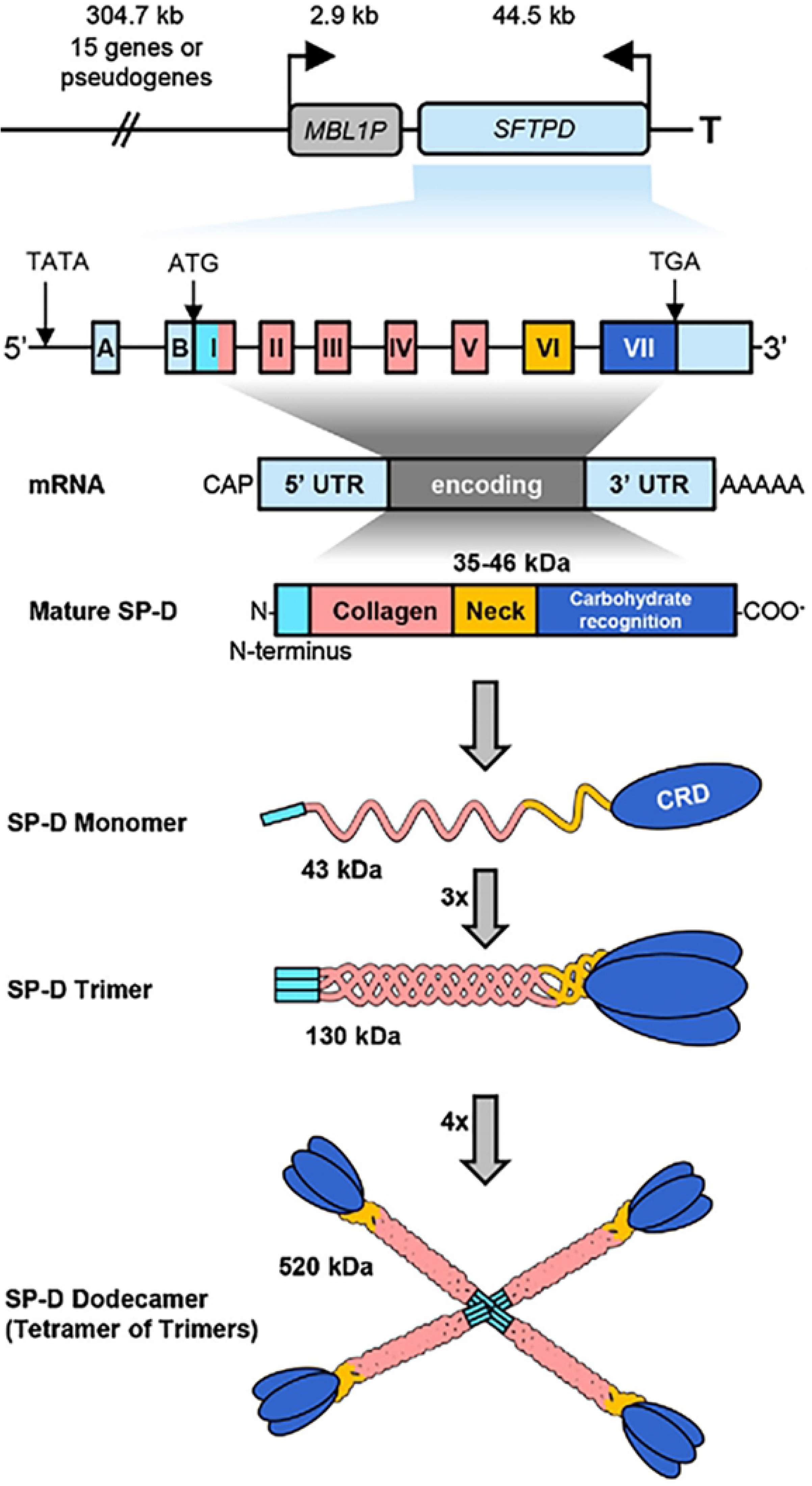
Schematic of gene map, transcription, and translational products and final protein structure of SP-D. The genes for SP-D are on chromosome 10. The mRNA is flanked by 5′ untranslated regions (UTR) and 3′ UTR containing a poly A tail. The UTRs are depicted in blue for Sftpd. There are two exons in the 5′UTR of SP-D (A, B). The four protein coding domains are named I-VII. The neck (VI) and collagen (I-VI) domains are shown in identical colors in the translated proteins. The carbohydrate recognitions domain (CRD, VII) and the cysteine-rich N-terminuses (I) are SP-D-specific. The final arrangement of protein chains is depicted with the classical cross of arms for SP-D. Modified from Vieira *et. al,* “Structure, genetics and function of the pulmonary associated surfactant proteins A and D: The extra-pulmonary role of these C type lectins”, Ann Anat. 2017

#### Immunohistochemistry (IHC)

Enucleated whole eyes were placed in fixative (PreFer; Anatech, Ltd., Battlecreek, MI, USA) for 30 minutes at room temperature (RT). Lung tissue was kept in the fixative solution until complete fixation was indicated by precipitation to the bottom of the container. Tissues were then treated with 70 % ethanol, embedded in paraffin, and sections were prepared on glass slides. Tissue sections were deparaffinized and blocked in 10% horse serum in PBS-0.1% Triton for 2 h. Sections were then incubated in the following primary antibodies overnight at 4 °C: rabbit polyclonal anti-SP-D (1:100 dilution; Antibodies-online Inc., St. Atlanta, GA, USA); rat anti-CD31 for endothelial cells (1:200 dilution; Dianova GmbH, Hamburg, Germany). Sections were incubated with Alexa Fluor 488- and 594-conjugated secondary antibodies (Invitrogen, Grand Island, NY, USA) at RT for 2 h. Sections were examined by confocal microscopy (SP2 model confocal microscope; Leica Microsystems GmbH, Buffalo Grove, IL, USA). All images shown are maximum projections from z-stacks through the entire tissue section. Primary antibody omission controls were performed for all antibodies (data not shown).

#### Western Blot for SP-D Protein Detection in the Mouse Retina

Dissected whole retina and lung tissue (positive controls) homogenates were prepared by the addition of 100 to 150 μL of lysis buffer (Invitrogen, Grand Island, NY, USA) with protease inhibitor cocktail (Millipore, Billerica, MA, USA), respectively. Tissue samples were sonicated and centrifuged, and the supernatant was collected. Total protein concentration was assessed by bicinchoninic acid (BCA) assay using a Pierce BCA Protein Assay Kit accordingly to the manufacturer’s instructions (Thermo Scientific; Lenexa, KS, USA). Equal volumes of 30 μg total protein were denatured by adding Sample Loading Buffers (BIO-RAD Laboratories; Hercules, CA, USA). Denatured protein samples were loaded onto 4-20% Tris-Glycine Gel Novex WedgeWell (Invitrogen; Grand Island, NY, USA) for gel electrophoresis. Protein from the gel was then transferred to a BioTrace Nitrocellulose transfer membrane with 0.2-μm pores (PALL laboratory; Port Washington, NY, USA). The membrane was blocked with 5% skim milk powder in tris-buffered saline (TBST). Membranes were incubated with primary antibodies at 4 °C overnight. The following primary antibodies were used: goat anti-SP-D from Santa Cruz (1:250 dilution; Dallas, TX, USA) and R&D systems (1:250 dilution; Minneapolis, MN, USA), and rabbit anti-SP-D from Antibodies-online, Inc. (1:250 dilution; Atlanta, GA, USA). Secondary antibodies, donkey anti-goat horseradish peroxidase (HRP) from Abcam (1:500 dilution; Cambridge, MA, USA) and donkey anti-rabbit from Invitrogen (1:500 dilution; Grand Island, NY, USA), were incubated at RT for 2 h and covered with foil. Membrane images were obtained using SuperSignal West Dura Extended Duration Substrate (Thermo Scientific; Lenexa, KS, USA) and Kodak Image Station 2000MM (Rochester, NY, USA).

#### Enzyme-Linked Immunosorbent Assay (ELISA) for SP-D Protein Quantification

Retina and lung homogenates were prepared as described above for western blotting. The Mouse SP-D Immunoassay Quantikine ELISA kit from R&D system (Minneapolis, MN, USA) was used according to the manufacturer’s directions. Briefly, the microwells of a 96-well plate were coated with diluted, purified anti-mouse SP-D monoclonal antibody. The wells were washed, and nonspecific sites were blocked. Diluted, purified mouse SP-D standards (0.625–40 ng/mL) and retinal lysates were added to the antibody-coated wells, and the plate was incubated for 2 h at RT. The plate was washed and incubated with a horseradish peroxidase-conjugated anti-mouse polyclonal SP-D antibody. Color reagents tetramethylbenzidine substrate solution and stabilized hydrogen peroxidase were added. The reaction was stopped by adding hydrochloric acid. The color change was measured by reading it at 450 nm, with the correction wavelength set at 540 nm or 570 nm using a FLUOstar Omega multi-mode microplate reader (BMG LABTECH, Cary, NC, USA).

### UP-REGULATION OF SP-D EXPRESSION IN THE MOUSE RETINA

After mouse retinas were examined at baseline, we sought to determine whether the expression of SP-D mRNA and protein could be elicited by inflammatory stress after intravitreal injections with the toll-like receptor ligands (TLR), TLR-2 and TLR-4, or after exposure to hyperoxia.

#### Intravitreal Injection of TLR-2 and TLR-4 Ligands

Six-week-old WT mice were anesthetized by intraperitoneal injection of ketamine/xylazine (100:10 mg/kg). Animals received 1 μg of the TLR-2 ligand Pam3Cys-Ser-(Lys)4 trihydrochloride (Pam3Cys) (Invivogen, San Diego, CA, USA), 1 μg of the TLR-4 ligand LPS (Sigma-Aldrich Corp., St. Louis, MO, USA), or control phosphate-buffered saline (PBS) in a total volume of 1 μL PBS vehicle. Injections were performed intravitreally using a 36-gauge needle mounted on a 10-μL syringe (Hamilton Co., Reno, NV, USA). The tip of the needle was inserted under the guidance of a dissecting microscope (Wild M650 model; Leica, Bannockburn, IL, USA) through the dorsal limbus of the right eye. The animals were euthanized at 6, 12, 24, and 48 hours after the injections. Whole-retina homogenates were used for RNA extractions and mRNA PCR, as described above.

#### Expression of SP-D in OIR

OIR was induced in WT and SP-D ^−/−^ mice by using a previously published technique (19). Briefly, all newborn mouse pups were maintained in room air (RA) with their dams until P7. At P7, pups (n=6) and the dams were placed in a poly(methyl methacrylate) (Plexiglas) chamber and exposed to 75% oxygen, using the Oxycycler C42 system (Biospherix, Lacuna, NY, USA). The second set of pups (n=6) were kept in room air to serve as controls. The dams were replaced every 48 hours with healthy dams, as adult mice do not tolerate prolonged hyperoxia. At P12, the pups and dams in the oxygen chamber were returned to room air and maintained there until P17. Whole retinas from WT animals were studied at three time-points for expression of SP-D mRNA: a) at P7 before oxygen exposure (baseline), b) at P12 after hyperoxia (vaso-obliterative [VO] phase), and c) P17 at the time of completion of OIR (neovascular [NV] phase).

#### Retinal Flat-mounts and Imaging

To examine the effect of the absence of SP-D on retinal vascular phenotype at baseline, after inflammation, and in OIR, mouse pups were euthanized, and their retinas were harvested and incubated with anti-CD31 antibody, followed by flat-mounting. Total retinal area, area of VO, and area of NV were analyzed using a well-established and standardized method (20). Briefly, Adobe Photoshop was used, and the area of vascular growth was outlined and quantified to measure the total retinal area, the area of VO, and the area of NV. NV was expressed as a percentage of the total retinal area. Six animals were included in each group, which had been determined by power analysis to be able to detect a 30% difference in NV with a b error of 0.2 and an area of 0.05. The mean areas of NV on the flat-mounts were compared with Student’s *t*-test. A P-value of < 0.05 was considered significant.

### MASS SPECTROMETRY (MS) FOR THE IDENTIFICATION OF SP-D PROTEIN

For final confirmation of SP-D expression, MS was performed on retinas from WT C57BL/6J mouse pups. WT and SP-D^−/−^ retina and control lung samples were prepared based on the standard targeted proteomics approach. Sixty μg of total protein each sample was taken and 8 pmol BSA added as an internal standard. The proteins were precipitated with acetone. The dried protein pellet was reconstituted in Laemmli sample buffer, and 20 μL (20 μg total protein) was run into a short (1.5cm) SDS-PAGE gel. The gel was fixed and stained. Each sample was cut from the gel as the entire lane and divided into smaller pieces. The gel pieces were washed to remove the Coomassie blue, then reduced, alkylated, and digested overnight with trypsin. The mixture of peptides was extracted from the gel, evaporated to dryness in a SpeedVac, and reconstituted in 150 μL 1% acetic acid for analysis.

Protein analyses were carried out on a QEx quadrupole-orbitrap mass spectrometry system. The HPLC was an Ultimate 3000 nanoflow system with a 10cm × 75μm i.d. C18 reversed-phase capillary column. Five-μL aliquots were injected, and the peptide eluted with a 60-min gradient of acetonitrile in 0.1% formic acid. The mass spectrometer was operated in the parallel reaction monitoring mode (PRM). The method was developed to initially target a total of seven peptides based on a combination of the protein sequence and information in the PeptideAtlas database. Data were analyzed using the program SkyLine to find and integrate the appropriate chromatographic peaks. These analyses showed that four of the possible peptides were detectable (Table 2).

**Table 2.** Peptides analyzed by Mass Spectrometry.

| Sftpd peptides | Domain location | SP-D specificity | Tissue of expression |
| --- | --- | --- | --- |
| AALFPDGR | Neck domain | Specific | Only WT lung |
| GENGSAGEPGPK | Collagen domain | Not specific | WT & KO - lung & retina |
| GESGLPDSAALR | Collagen domain | Not specific | WT & KO lung & retina |
| LEVAFSHYQK | Neck domain | Specific | Only WT -lung |
| GEVGAPGMQGSTGAK | Collagen domain | Not specific | WT & KO lung & retina |
| AAFLSMTDVGTEGK | CRD domain | Exclusively SP-D | Only WT lung |
| SATENAAIQQLITAHNK | CRD domain | Exclusively SP-D | Only WT lung |

The peptides highlighted in gray are specific for SP-D and were only found in the WT lung. The non-specific peptides were identified in lung and retina in the WT and KO mice. Please refer to supplemental material for more details.

## Results

### Is SP-D gene expression detected in the mouse retina?

Utilizing RT-PCR, SP-D mRNA was detected with positive bands at 430 bp in the WT lung tissue, which were absent from the SP-D^−/−^ lung. In the WT retina from healthy animals, the expected band at 430 bp was not detected (Figure 6). Similarly, in the postnatal developmental retina sample, SP-D mRNA was not present in the retina (Figure 7b).

The absence of gene expression at baseline in WT mice prompted us to evaluate whether expression could be induced in a stressful pathological state. First, we tested retinal tissue from WT pups exposed to the OIR model at both P12 (VO) and P17 (NV) (Figure 8). Exposure to oxygen did not induce the expression of SP-D mRNA. We also tested retinas exposed to Pam3cys (TLR-2 ligand) and LPS (TLR-4 ligand) by intravitreal injection (Figure 7c). These insults did not induce the expression of SP-D mRNA.

### Expression of SP-D Protein in the Mouse Retina

Initially, our primary antibody was tested in WT C57BL/6J adult mouse lung tissue. Expression of SP-D, according to IHC results, is characterized by a speckled appearance in the cells lining the alveolar spaces in the color green due to secondary antibody Alexa-Fluor 488 (Figure 2a). SP-D expression was seen in retinal cross-sections in adult mouse retinas. We then proceeded with an analysis of cross-sections of the retina at early developmental time points (Figure 2a and 2h). However, our antibody strategy gave expected results only at P0 (Figure 2b). At older ages from P2 to P14, the detection of SP-D was decreased (Figure 2c - f).

**Figure 2.**
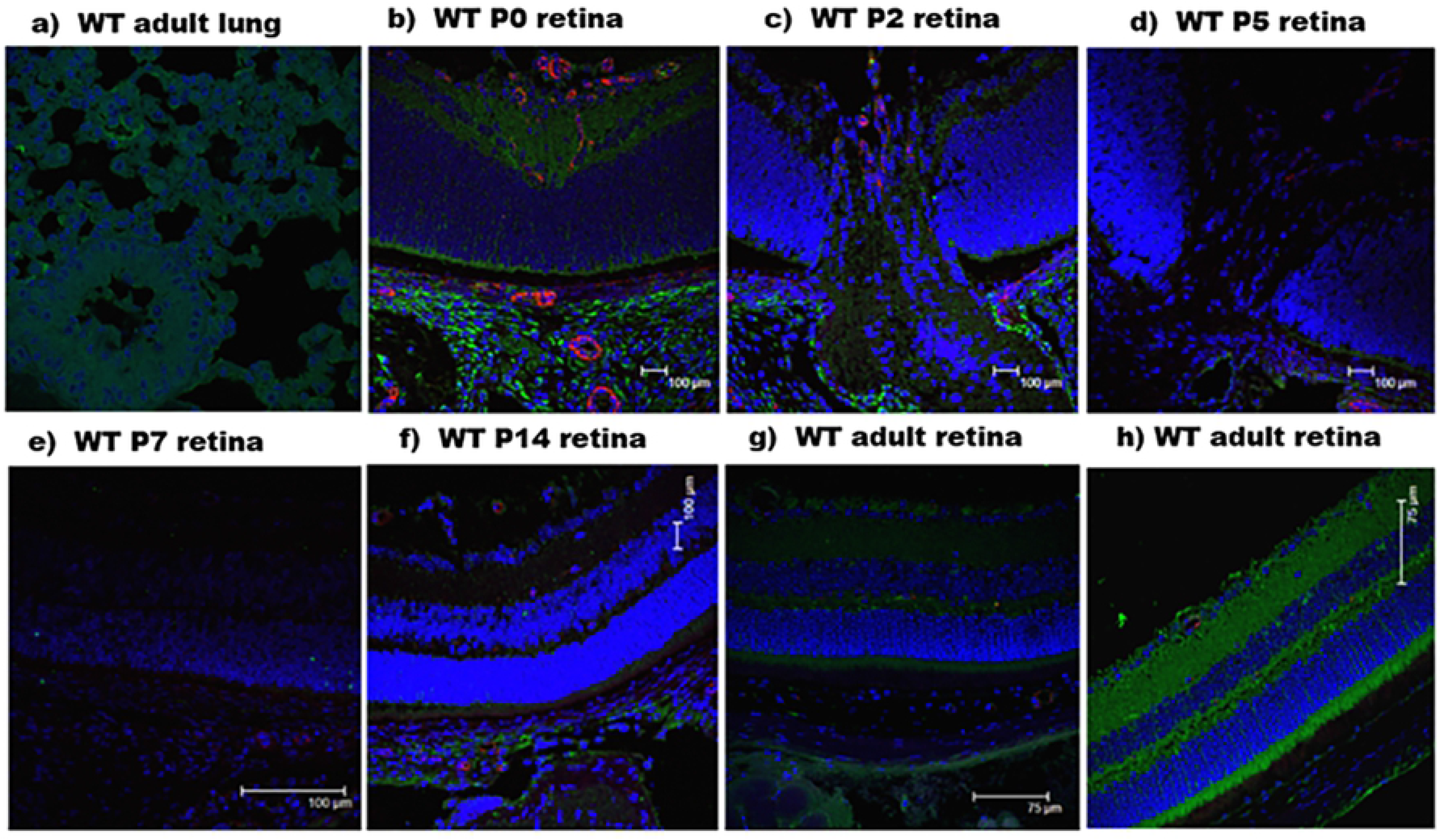
C57BL/6J wild-type (WT) mouse immunohistochemistry. Green color represents SP-D localization. Blue indicates cellular nuclei stained by DAPI. Red indicates vessel stained with anti-CD31. a) Adult lung tissue. b), c), d), e), f), g) show staining of the central retina at developmental stages from P0 to adult mouse. h) adult peripheral retina.

Similarly, the western blot of the homogenized retina from WT C57BL/6J mouse did not show consistent bands for SP-D protein. The R&D systems antibody gave a 45 kDa band for WT lung tissue, as expected. However, the WT retina tissue did not generate any specific band. For SP-D ^−/−^ animals, the lung tissue was negative, and the retina tissue gave a band of 69 kDa. The primary antibody (Antibodies-online, Inc.) gave multiple positive bands in the WT lung tissue and SP-D ^−/−^ lung and retina. The WT retina did not produce any strong bands (Figure 3). This led us to conclude that readily available SP-D antibodies are not reliable in the detection of protein in tissues other than the lung.

**Figure 3.**
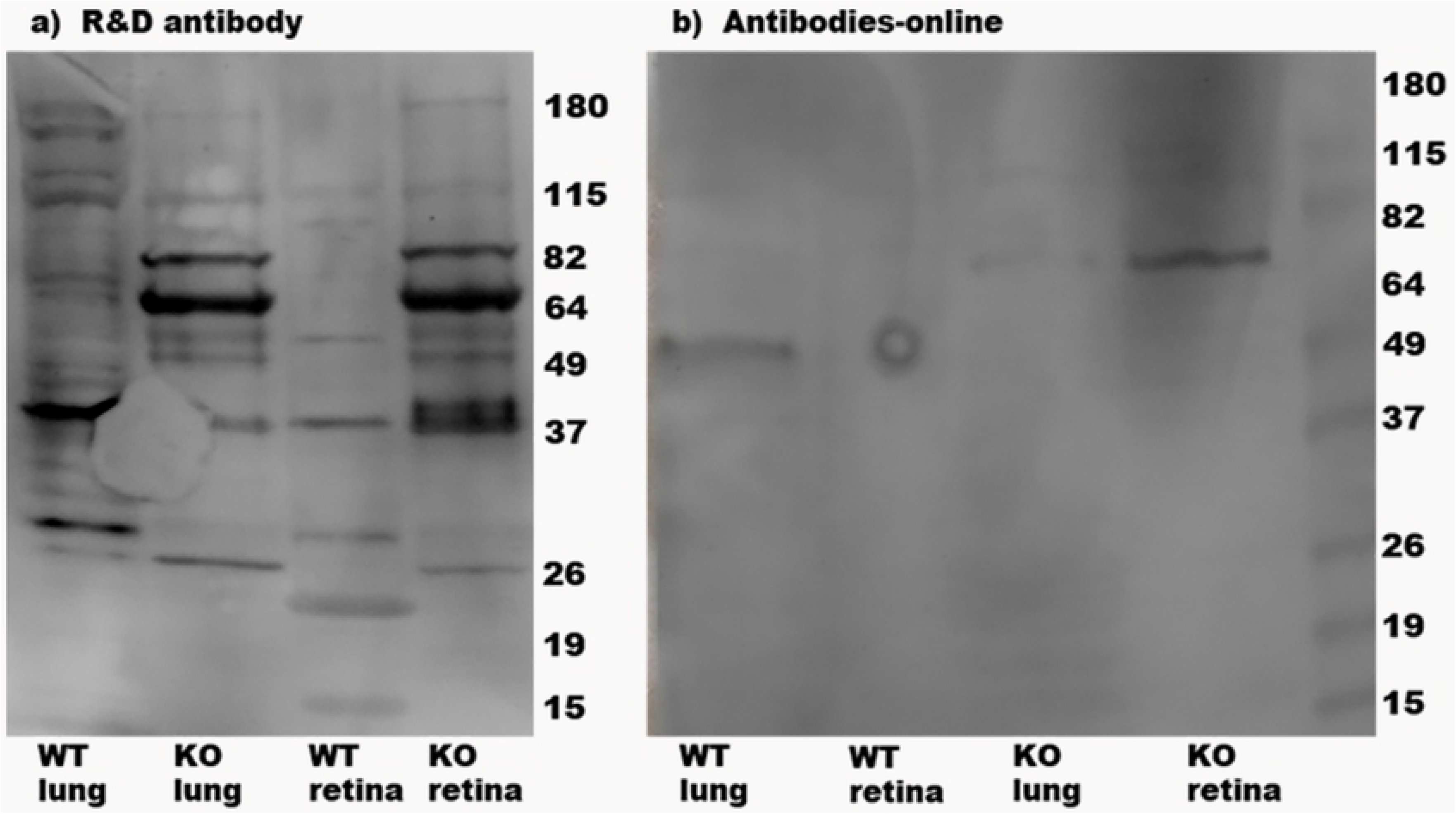
Western blot for SP-D in lung and retina for WT and SP-D^−/−^ mouse.

For further confirmation, SP-D ELISA was performed. It was also negative for SP-D in the adult WT retina. The adult WT lung sample had approximately 3 ng/mg of total protein. However, the WT adult retina had an undetectable amount of SP-D.

Since methods to detect SP-D with commercially available antibodies were not successful, we then proceeded to protein mass spectrometry. We sent retina and lung tissues from WT and SP-D ^−/−^ mice for peptide analysis. SP-D peptides identified in the WT lung and not in the SP-D ^−/−^ lung are reported in Table 2. These peptides belong to the CRD domain and the neck domain, which are the collagen domains with no similarities to other collectin proteins (Figures 4 and 5). The peptides that were not present in the SP-D ^−/−^ lung and retina tissue were also not present in the WT retina tissue, demonstrating that it is unlikely that SP-D was present in the retina.

**Figure 4.**
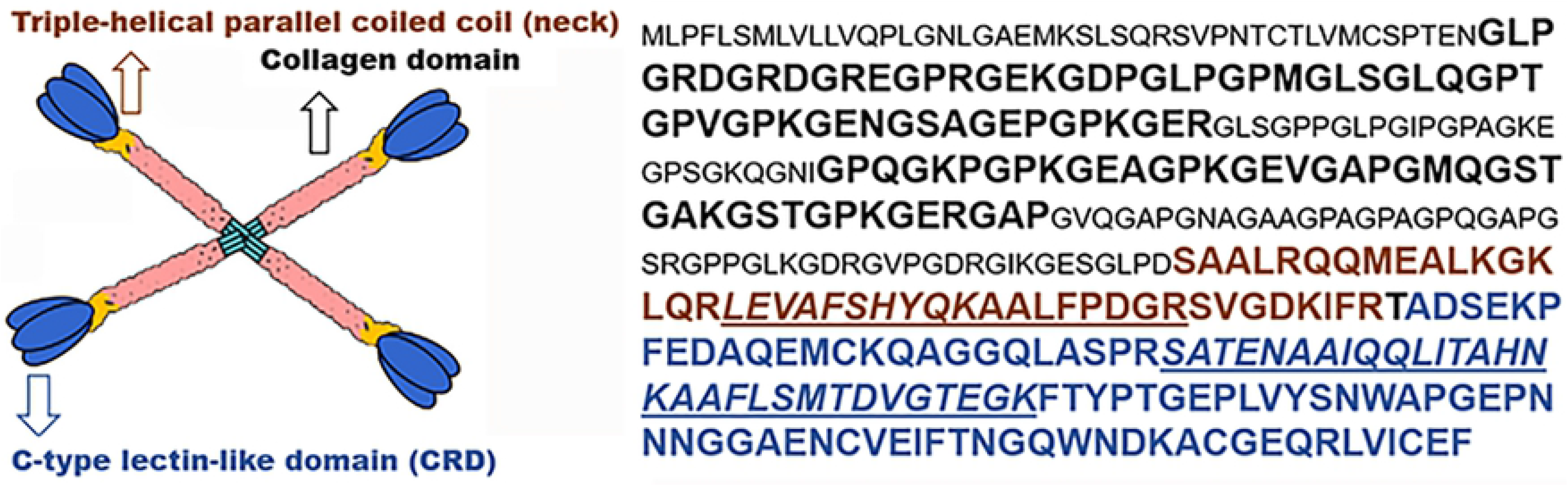
Peptide sequence in mature SP-D. Bold black represents peptides from the collagen domain, most similar to other collectins. The neck domain (brown) and CRD (blue) have different nucleotide sequences from other collectins. Underscored amino-acids represent the peptides sequence present only in the WT lung, but not in the SP-D^−/−^ lung and WT retina, according to mass spectrometry.

**Figure 5.**
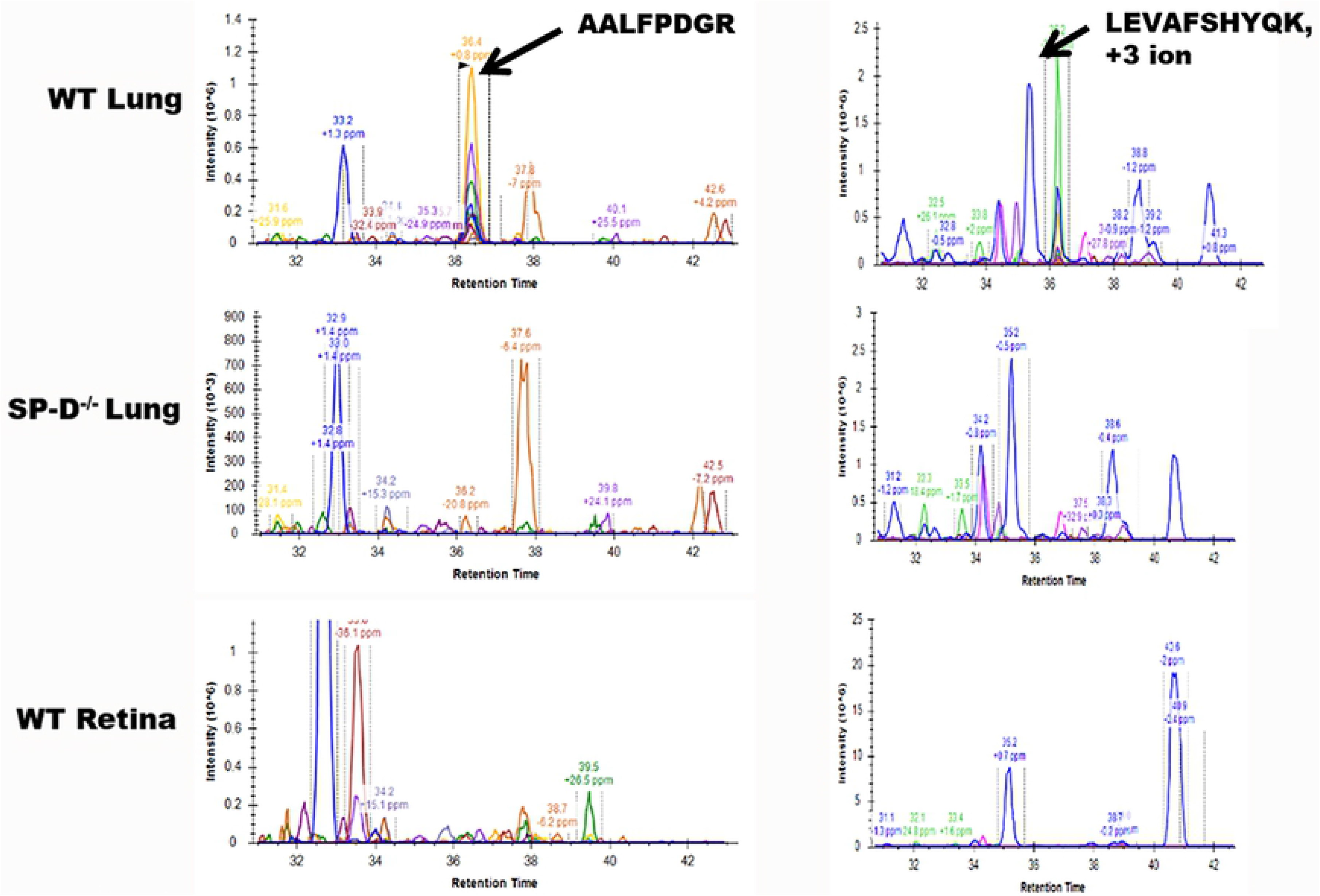
Mass spectrometry results. Two peptide sequence of mature SP-D, “AALFPDGR” and “LEVAFSHYQK”, of the neck domain has a peptide peak in the WT lung, but not in the SP-D^−/−^ lung or WT retina.

**Figure 6.**
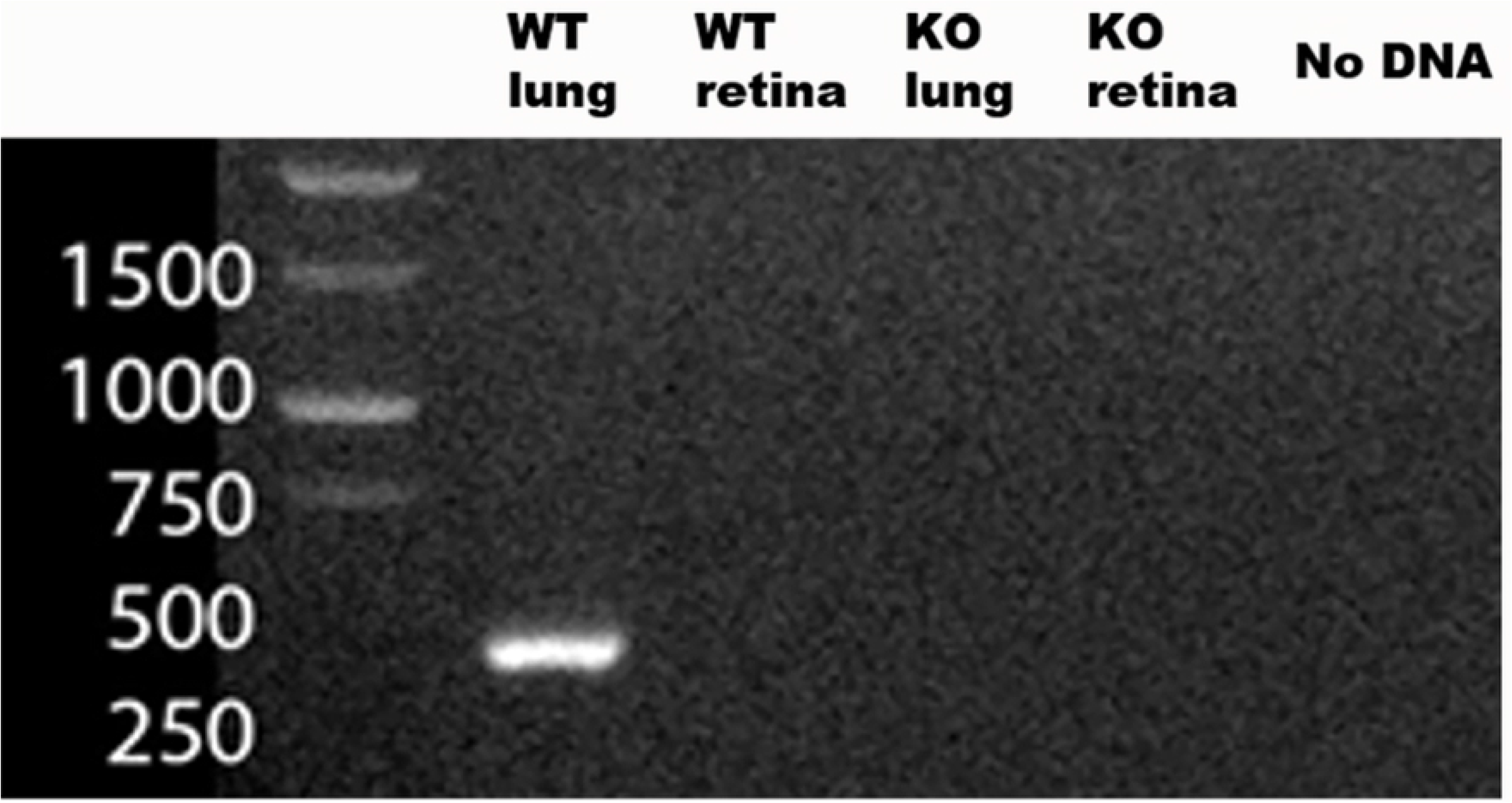
mRNA PCR for SP-D (430 Kb pair). Positive for WT lung (positive control), negative for SP-D^−/−^ lung (negative control). WT retina and SP-D^−/−^ retina were negative.

**Figure 7.**
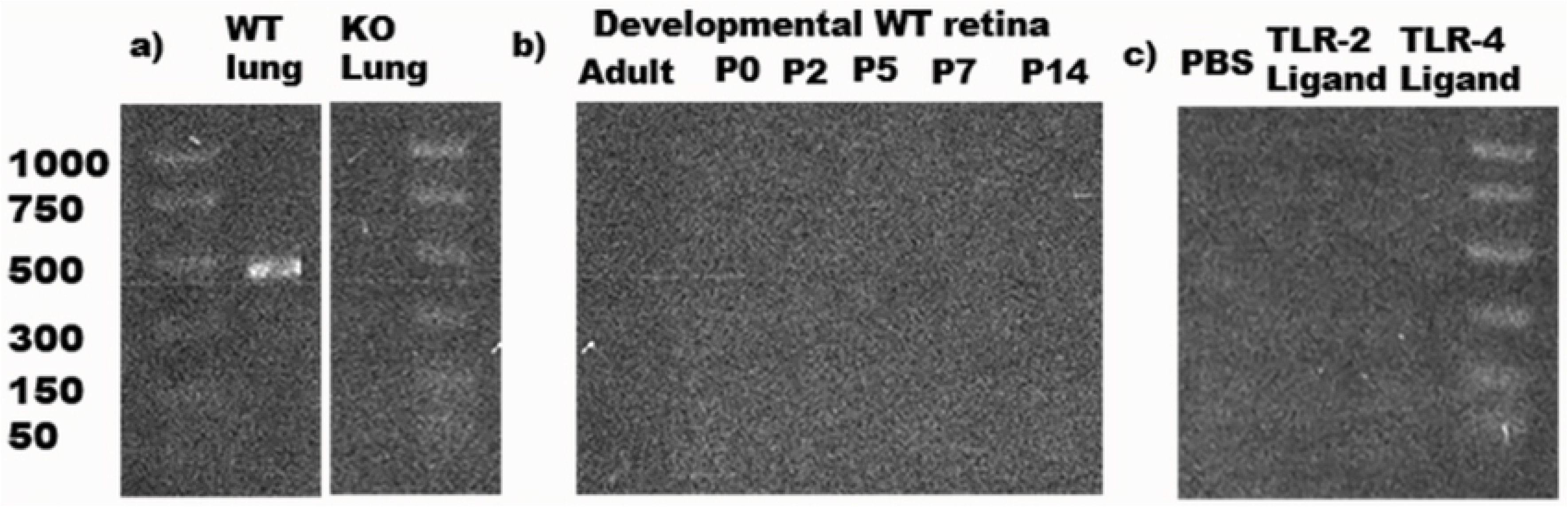
mRNA PCR for SP-D. a) WT lung and KO lung positive and negative controls; b) developmental retina from P0 to adult mouse; c) intravitreous injection of TLR-2, TLR-4 ligands, and PBS (negative control). SP-D mRNA at 430 kb.

**Figure 8.**
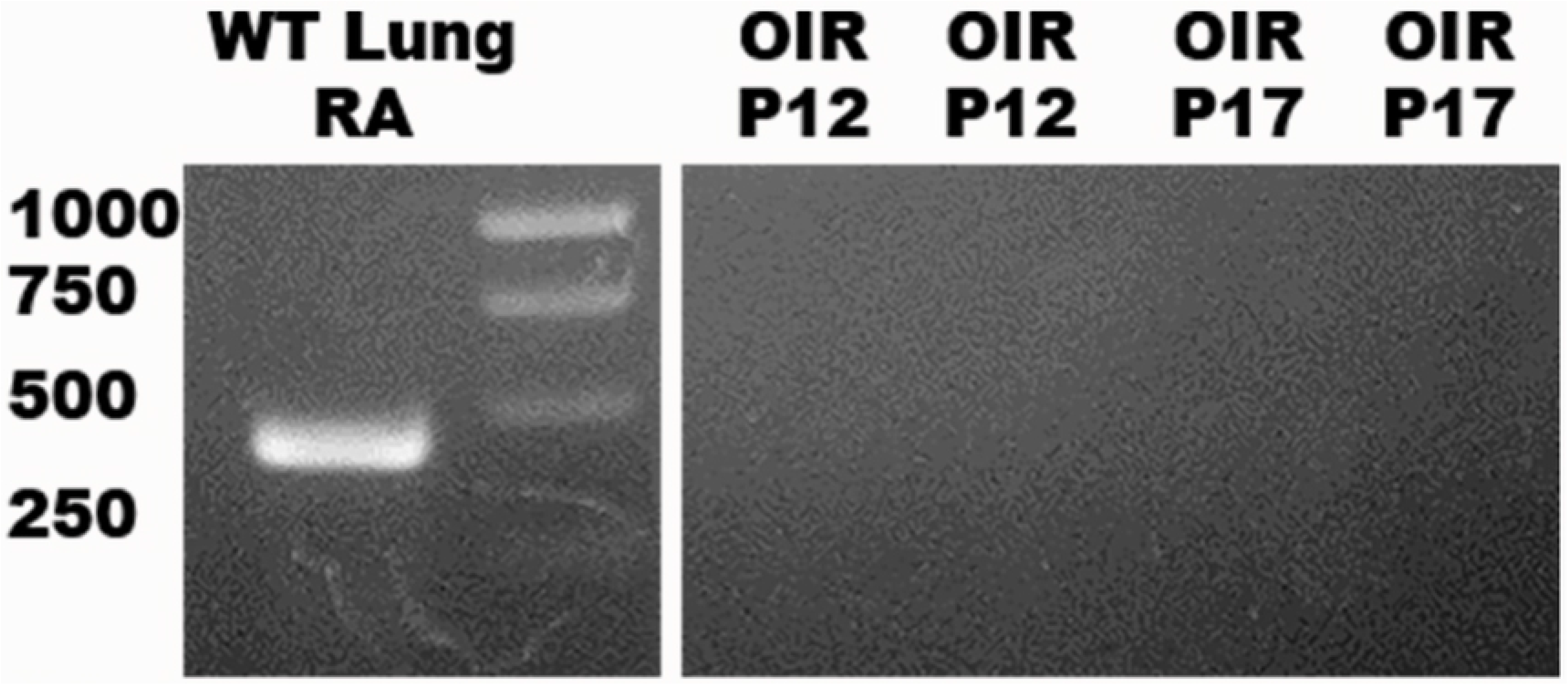
mRNA PCR. WT lung on RA (positive control) generated 430 kb. Two samples of OIR P12 and P17 were negative.

### Does SP-D Impact the Vascular Phenotype in the Mouse Oxygen-Induced Retinopathy Model?

WT and SP-D ^−/−^ mice were exposed to hyperoxia to induce VO and NV in the OIR model. WT mice had an NV ratio of 21%, while SP-D ^−/−^ mice had 16%. While there was a modest decrease in NV of 5% in SP-D ^−/−^ mouse retinas, this difference was not statistically significant (p-value 0.3531). Vaso-obliteration ration was 6% in both WT and SP-D ^−/−^ mouse retinas (p-value of 0.9957). WT mouse retinas and SP-D ^−/−^ in RA that were not exposed to OIR had no NV, as expected (Figure 9).

**Figure 9.**
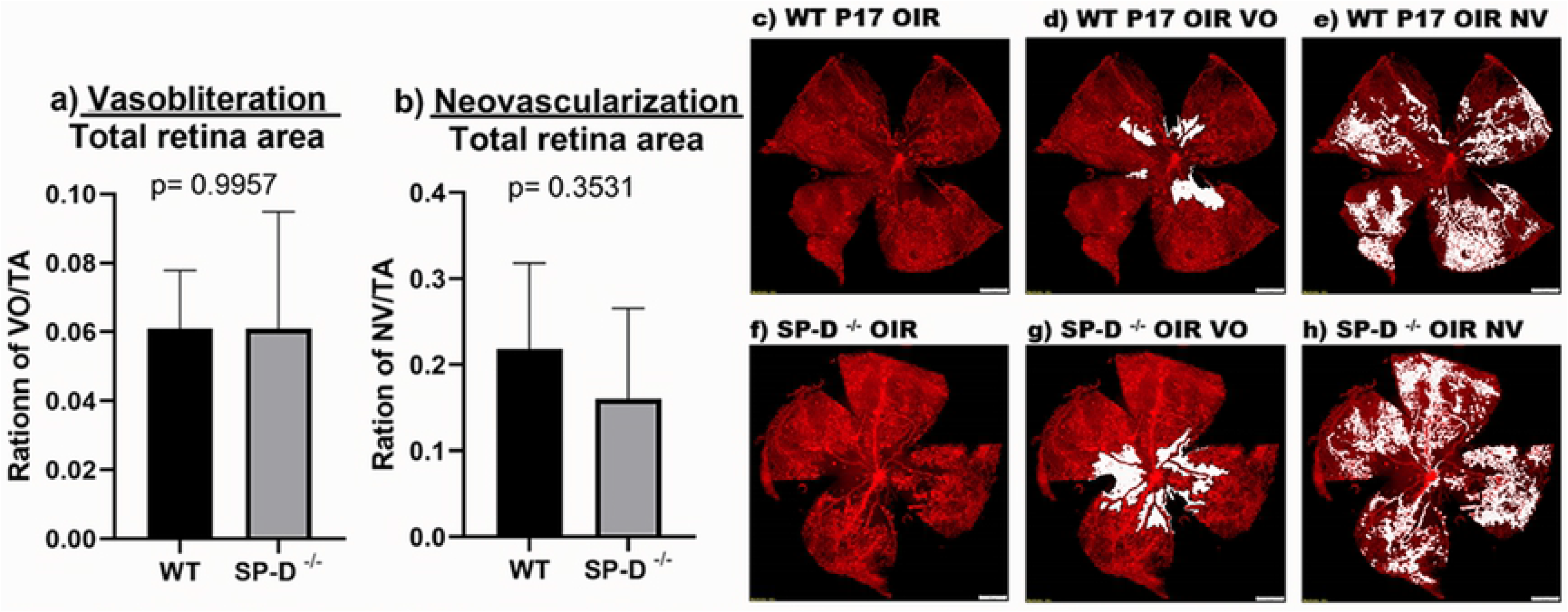
a) Vasobliteration and b) Neovascularization ration in WT and SP-D^−/−^ C57 pups at P17 after oxygen-induced retinopathy (OIR); N=6 per group. (c, d, e) Example of retina flat mount of C57BL/6J wild-type animal after OIR. (f, g, h) Similar example of retina flat mount of SP-D^−/−^ mouse. Vasobliteration (VO). Neovascularization (NV)

## Discussion

SP-D is an important c-type lectin that was initially thought to be primarily associated with pulmonary tissue, but has been reported to be present in a variety of organ systems in mice and humans (13). This study is the first to investigate the presence of SP-D in the mouse retina. We have demonstrated that SP-D is not present in the C57BL/6J mouse retina at baseline, nor up-regulated by inflammatory or oxidative stress. The results were confirmed by gene expression and proteomic analysis.

Retinal vascular disease (ROP) is a significant long-term complication for preterm low birth weight infants, and the leading cause of acquired childhood blindness in the U.S. (21, 22). The pathophysiology of ROP is multifactorial, but inflammation is one of the major contributors to the disease (23–25). Vascular endothelial growth factor (VEGF) is an essential factor in the abnormal neovascularization seen in ROP (26). Several studies have shown that Toll-Like Receptors 2 and 4 up-regulate VEGF in lung tissue after exposure to hyperoxia (27). Also, VEGF will up-regulate SP-D production in type II pneumocytes in the lung (28). Furthermore, there is a direct correlation of TLR-4, causing retinal ischemia/reperfusion injury (29). We therefore speculated that a variety of stressors and inflammatory signaling pathways may lead to the expression of SP-D in the retina, and that SP-D could participate in the abnormal vascularization seen in ROP. Since SP-D has been identified in multiple tissues other than the lung, including brain tissue (13), the presence of SP-D in retina seemed plausible. Furthermore, SP-A, an analogous c-type lectin protein, is present in the retina, is up-regulated by TLR-2 and 4 ligands and in the OIR mouse model, and affects both physiological (unpublished data) and pathological retinal vascularization (14).

### SP-D, Inflammation, and the Prematurely Born Infant

Surfactant Protein D belongs to a superfamily of c-type lectins with multiple immunological functions. One of the defining characteristics of SP-D is the CRD affinity for inositol, maltose, and glucose, and a poor affinity for sugar moieties of galactose and sialic acid in invertebrate animals (30). These differences in carbohydrate affinity are important in identification of pathogens in the airway and other tissues, but not self-sugar moieties. It also plays a crucial role in the ability of SP-D to opsonize bacteria and viruses (31–33) and direct the killing of pathogens (34). SP-D has been shown to bind to multiple receptors and acceptors, which may result in either pro- or anti-inflammatory activity. Scavenger receptor cysteine-rich (SRCR) (35), defensins (36–38), calreticulin (39), TLR-2 and 4 (10), LPS (TLR-4 ligand) (40), and CD14 (co-receptor of TLR-4) (41) are among the acceptors and receptors important in the innate immunity with which SP-D interacts. Even though SP-D participates in adaptive immunity (42), the main function of SP-D correlates with innate immunity. Taken together, these characteristics define the importance of the SP-D in the medical specialty of neonatology, in which subjects will have an inflammatory response-related function of the innate immune system.

### SP-D Expression in the Mouse Retina

The C57BL/6J mouse was chosen as the subject for our investigations as its retinal vascular development is similar to that of a preterm newborn delivered at approximately 24 weeks’ gestational age (43). Initial pilot and feasibility experiments involved the examination of developmental immunohistochemistry, since cross-sectional retinal tissue can be easily obtained at postnatal ages P0, P2, P5, P7, P14, and P42-P48 (adult). In addition, the necessary primary antibodies for IHC were readily commercially available. Lung tissue of WT animals was used as a positive control for our antibodies. The tested antibodies localized to the airway lining of pulmonary alveoli, an area with very high expression of SP-D. When retinal tissue was tested, IHC showed strong staining by the antibody, with localization in the optic nerve, optic disk, and choroid tissue, areas known to have greater vascularization in neonatal mice, at P0. At ages P5, P7, and P14, there was a significant decrease in antibody binding in the retina tissue and optic nerve. However, when appropriate negative controls (tissue from SP-D ^−/−^ mice) were evaluated, there were inconsistent results in retinal tissue. Further evaluation with ELISA and western blot analysis was therefore performed. ELISA for SP-D showed a high concentration of SP-D protein in WT lung control. However, we were never able to measure SP-D in the retina tissue of WT adult mice. In our western blot, we had a similar issue with our commercially acquired antibodies. We detected SP-D in WT lung tissue, but it did not give a band of 45 kDa in WT retina tissue. For this reason, testing with a variety of different commercially available antibodies was performed. However, these antibodies either showed no expression in retina protein extract, or they produced multiple bands at different molecular weights. We speculate that since the commercially available antibodies are generated for pulmonary SP-D, retinal protein may have a different post-translational modification or haplotype due to gene polymorphism (44, 45). The primary antibodies with positive results were polyclonally derived, making it impossible to determine which of the clones would be expected to bind to the CRD domain of the monomeric structure of the protein after denaturation. As described earlier, the CRD domain has an amino acid sequence distinct from that of other collectin proteins and has minimal variants among haplotypes.

Due to the uncertainty of the affinity from commercially available antibodies to detect expected retinal SP-D, WT retinal tissue was sent for proteomic mass spectrometry (MS). Mass spectrometry has the advantage of analyzing peptide sequences according to molecular weight and charge without a need to use any other biological support to identify proteins or peptides (46). However, the peptide libraries for WT mouse retina, PeptideAtlas database, had no reported information for SP-D. For appropriate comparison, adult WT lung and SP-D ^−/−^ lung was sent to compare major peptides that would be present in the WT tissue, but not in the gene-deleted tissue. In this manner, peptides from proteins other than SP-D would not interfere in the search for SP-D in the WT retina. The peptides absent from the SP-D ^−/−^ lung were also absent from the WT retina. Of these four peptides, two were from the CRD domain and two were from the neck region, which are the regions with the lowest variation among haplotypes and minimal similarities to other collectin proteins.

Bhatti *et al*. have previously reported expression of SP-A mRNA and protein in whole murine retinal tissue, as well as in cultured human MIO-M1 and Müller cells (14). We, therefore, hypothesized that Müller cells would similarly express SP-D in the mouse retina. Previous proteomic analysis by MS of murine retinas of Müller cell-enriched and -depleted samples did not identify SP-A or SP-D (47). However, the analysis was guided by the concentration of peptides, so we could argue that SP-D was not demonstrated in MS of the WT retina due to low concentration in retinal Müller cells.

In summary, our proteomic investigation of the possible presence of SP-D in the mouse retina showed that commercially available antibodies to pulmonary SP-D did not detect SP-D in the retina. Mass spectrometry did not detect important peptides that define SP-D in the mouse retina.

### Genomic Verification of SP-D in Mouse Retina

To supplement our proteomic results that SP-D is absent from the WT mouse retina, we evaluated the presence of intermediary pathway product SP-D mRNA, with WT lung used as a positive control. The results of the preliminary gel agreed with our proteomic results. PCR analysis showed that the WT lung was positive and the WT retina and SP-D ^−/−^ lungs were negative for SP-D mRNA. No SP-D mRNA was detected at the ages of P0, P2, P5, P7, P14, and P42-48.

Considering that SP-D is an immunoregulatory protein, it was possible that SP-D would be only minimally expressed at baseline and up-regulated in the retina once exposed to noxious stimuli as TLR-2 and TLR-4 ligands, which are known to up-regulate SP-D expression in lung tissue. As inflammation is a significant contributor to ROP and SP-A is up-regulated after intravitreal injection of Pam3Cys (TLR-2 ligand) and LPS (TLR-4 ligand), we decided to perform similar experiments in the WT mouse retina. However, no mRNA SP-D was detected in the retinas exposed to TLR 2 and 4 ligands.

Since our ultimate goal was to investigate possible effects of SP-D in ROP, we finally proceeded to study the impact of hyperoxia/hypoxia as a potential promoter of SP-D transcription in the WT retina. WT pups were exposed to the OIR model, which mimics the environment and pathological process endured by premature infants (19, 48). Retinas were harvested after five days of exposure to 75% hyperoxia at P12, at which point the retinal vasculature undergoes vaso-obliteration, as it does after relative hypoxia at P17, which would induce neovascularization similar to ROP. As discussed earlier, VEGF is the main signaling pathway for abnormal vessel development in this situation. Since VEGF up-regulates SP-D in lung tissues, hypoxia and hyperoxia could be additional noxious stimuli to potentially cause transcription of SP-D in the mouse retina. Once again, no SP-D mRNA was detected in any of the retinas at P12 and P17. In summary, SP-D is not transcribed in the WT mice, even after application of noxious stimuli known to cause up-regulation of this protein in other tissues.

### Retina phenotype of SP-D ^−/−^ mouse

At baseline and after noxious stimuli, we did not demonstrate the presence of SP-D in the retina at the molecular level. However, not all interactions of SP-D were postulated in this manuscript; this protein can have many relationships with different receptors and pathogens (13). In order to determine whether another possible pathway or receptor was impacted by the systemic absence of SP-D in the global knockout, retinal vascular phenotypes were compared between WT and SP-D mouse retinas after OIR. While there was a small decrease (5%) in retinal vascular disease, this difference was not significant. Therefore, while there may be a small contribution, the lack of systemic SP-D did not demonstrate a significant impact on the final phenotype in OIR.

## CONCLUSIONS

SP-D is not present in the WT C57BL/6J mouse retina. Proteomic analysis did not detect or measure SP-D from retina protein homogenate. The positive fluorescence in the preliminary IHC shows that the pulmonary SP-D antibody may be detecting other collectins in the retina tissue. Furthermore, the genomic investigation of SP-D mRNA demonstrated that SP-D is not being transcribed or generated by pathways known to up-regulate SP-D in lung tissue. Similarly, the absence of SP-D in the retina did not change the phenotype of vessel development in this tissue. From these findings, it is unlikely that SP-D participates in retinopathy of mouse retinas. While SP-A and SP-D are commonly co-expressed in a variety of tissues with a similar pro/anti-inflammatory signature, the mouse retina did not show such an association.

## ACKNOWLEDGEMENTS

The authors thank the Histology Core Facility and Imaging Core Facility; Mark Dittmar, Manager, Animal Facilities at the Dean McGee Eye Institute; and Mike Kinter at the Laboratory for Molecular Biology and Cytometry Research at the University of Oklahoma Health Sciences Center in Oklahoma City for assistance with mass spectrometry and proteomics services.

Supported by a U.S. National Institutes of Health P20 RR017703-10 Pilot Project Award and the 5P30EY021725 P30-Center Core Grant For Vision Research

